# Polar bodies serve as a landmark for anteroposterior axis formation in spiders

**DOI:** 10.1101/2024.09.30.615744

**Authors:** Ruixun Wang, Matthias Pechmann

## Abstract

The early embryogenesis of many spiders involves the formation of a radially symmetric germ disc. While the cells of the rim of this germ disc develops into anterior structures, the center of the disc will form the posteriorly located segment addition zone of the embryo. Therefore, germ disc formation sets the anterior-posterior (AP) body axis of spider embryos. The early spider egg is a spherical structure with no apparent asymmetry, and it is believed that germ disc formation is a stochastic process.

For this study, we have re-analyzed early spider embryogenesis and found a strong correlation of the position of the polar bodies and the formation of the germ disc. Our results suggest that germ disc formation in the common house spider *Parasteatoda tepidariorum* is not a stochastic but a pre-determined process. Furthermore, we provide evidence that this correlation might be conserved between araneomorph and mygalomorph spider species.

## Introduction

A great variety of mechanisms are involved in setting up the primary body axes in bilaterally symmetric animals. In many animals the anteroposterior (AP) as well as the dorsoventral (DV) body axes are established before or shortly after fertilization. Well-known examples are insect embryos that develop from a pre-patterned system that gets established already during oogenesis or amphibian embryos in which a rotation of the egg cortex is triggered by the sperm entry and is leading to a redistribution of maternally loaded components, which are required to initiate the AP as well as the DV body axes (e.g. Barresi and Gilbert 2020).

In many araneomorph spider species like *Parasteatoda tepidariorum* (syn. *Achaearanea tepidariorum* (Yoshida 2008)), early embryogenesis is characterized by the formation of a distinct germ disc in one hemisphere of the egg (e.g. Akiyama-Oda and Oda 2003, Yamazaki et al. 2005, Hilbrant et al. 2012). After the formation of a regular blastoderm, coordinated cell shape changes, changes in the quantity of yolk in certain cells and the onset of zygotic transcription are required to drive germ disc condensation in *P. tepidariorum* embryos (Pechmann 2016). The temporal and spatial expression analysis of certain marker genes revealed that the ectodermal cells that are located at the rim of the germ disc will develop into anterior structures of the developing embryo (e.g. Pechmann et al. 2009, Hemmi et al. 2018). In contrast, the ectodermal cells of the centre of the disc will end up in posterior regions of the developing embryo (e.g. McGregor et al. 2008, Hemmi et al. 2018, Schönauer et al. 2016). Clonal analysis supported these findings and demonstrated that germ disc formation is tightly linked to the formation of the anteroposterior body axis (Hemmi et al. 2018).

As spider eggs are spherical in shape, it so far is unclear if the initial placement of the germ disc is a stochastic or a predetermined process. Recently it was even stated: “During the initial developmental stages (i.e., stages 1 and 2), no morphological landmarks exist to indicate future body axes” (Oda and Akiyama-Oda 2020). However, previous studies have shown that in the common house spider *Paratsteatoda tepidariorum*, the polar bodies are present at stage 1 of embryogenesis (see Fig. 2b of Mittmann and Wolff 2012 and figure supplement 3 B’ of Pechmann et al. 2017). Mostly, polar bodies are the unused cellular by products of female meiosis that are developmentally suppressed and/or extruded from the egg (adapted quote from Peel and Averof 2010). However, the position of polar bodies often correlates with the formation of certain body axes. In the beetle *Tribolium castaneum* it was shown that the position of the polar bodies correlates with the formation of the DV body axis (Peel and Averof 2010, Lynch et al. 2010). Even in mammalian embryos like mice, the location of the polar bodies correlates with the formation of animal-vegetal body axis (reviewed in e.g. Beddington and Robertson 1999).

To see whether the initial placement of the germ disc shows any correlation with the position of the polar bodies, we established several new methods to visualize polar bodies in living and fixed spider embryos. Indeed, our analysis shows a clear correlation of germ disc formation and the early position of the polar bodies and suggests that germ disc formation in spiders is predetermined and not a stochastic process.

## Materials and Methods

### Spider husbandry and embryology

*Parasteatoda tepidariorum* and *Steatoda grossa* originated from the same cultures and were kept under the same conditions as described previously (Wang et al. 2023). A culture of *Ischnothele caudata* was newly established in our laboratory (Medina-Jiménez et al. 2024). Founder animals were purchased from German breeders. Adults and juveniles of *Ischnothele caudata* were kept in 19.5×19.5×11.5 BraPlast plastic containers at 22-25°C with a 12-hour day-night rhythm. According to the size, spiders were fed with *Drosophila melanogaster* or medium sized *Gryllus bimaculatus*. Embryos of *P. tepidariorum* and *S. grossa* were staged according to the published staging systems for *P. tepidariorum* (Mittmann and Wolff 2012). As the embryonic development is very similar, *Ischnothele caudata* was staged according to the published staging system for *Acanthoscurria geniculata* (Pechmann 2020).

Embryos were fixed with formaldehyde in a two-phase fixative solution as described previously (Pechmann et al. 2017). Formaldehyde fixed *P. tepidariorum* embryos were halved using tungsten needles.

Stage 1 *I. caudata* embryos were heat and formaldehyde fixed as it was described for *Gryllus* embryos (Pechmann et al. 2021).

Fixed embryos were counterstained with DAPI or SYTOX Green.

### Imaging and image analysis

Pictures were captured using an Axio Zoom.V16 (Zeiss; equipped with an AxioCam 506 color camera) and an AxioImager.Z2 (Zeiss; equipped with an Apotome.2 module and an AxioCam MRm camera). Helicon Focus and Zen2 software was used to create projections of the image stacks. Live imaging was carried out as described previously (Pechmann 2020). Tracking of nuclei and further processing of images and movies was executed using Fiji (Time stamper and MTrackJ plugins) and Photoshop CS5.1 (image labelling and adjustments for brightness and contrast). Drawings were created using Photoshop CS5.1 and Inkscape.

### Immunohistochemistry

After fixation, embryos were stepwise rehydrated, washed with PBST (3 × 15 min) and blocked in PBST containing 0,1 % BSA and 5 % goat serum (1 hr at RT). Embryos were incubated (o.n. at 4 °C) with a 1:1000 primary antibody solution (1 ml PBST/BSA/NGS [final BSA concentration 0,1 %, final NGS concentration 5 %] containing 1 μl of Phospho-Histone H3(Ser10) polyclonal antibody; Invitrogen; PA5-17869). Subsequently, the embryos were washed in PBST (3 × 15 min) and were blocked again in PBST containing 0,1 % BSA and 5 % goat serum (1 hr at RT). To detect the primary Phospho-Histone H3 antibody, a Goat anti-Rabbit IgG (H+L) Highly Cross-Adsorbed Secondary Antibody, Alexa Fluor™ 647Anti-Rabbit IgG, (Invitrogen; A-21245) was added to the blocking solution at a 1:1000 concentration. After incubating the secondary antibody for 2–3 hr (RT), excessive antibody was removed by washing the embryos several times in PBST (6 × 15 min; final washing step at 4 **°**C o.n.). Before imaging, embryos were co-stained with DAPI and mounted in 70 % glycerol.

### Whole mount *in situ* hybridization

A 2.362 bp fragment of *Pt-Delta* (AUGUSTUS number: aug3.g25248.t1 (Schwager et al. 2017)) was amplified using Pt-25248-F: 5’-CAT GCG GTG GTG TTA TTT AGT ATT ACT C-3’ and Pt-25248-R: 5’-CAA TTA CCT TTT GCT CGA TGC TAT ATT TG-3’ primer. The PCR product was cloned into Zero-Blunt-TOPO Vector (ThermoFisher). After cloning, M13F and M13R primer were used to generate a DNA template for the following synthesis of an antisense DIG-labelled RNA probe. *In situ* hybridization and false colour overlays of *in situ* hybridization images was performed as described previously (Pechmann et al. 2017).

### Fluorescent live imaging

SPY555-DNA (Spyrochrome) was dissolved in DMSO, mixed 1:1 with ddH_2_O and was injected to the yolk of stage 1 *P. tepidariorum* embryos. Microinjections were performed as described before (Pechmann 2016, Kanayama et al. 2010). Fluorescent live imaging was initiated around 1h after injection. Capped mRNA injections were performed as described before (Pechmann et al. 2017). To generate the construct for nuclear localized mCherry, EGFP was replaced by mCherry (see Figure 5—figure supplement 1 of Pechmann et al. 2017).

## Results

### Correlation between germ disc formation and the location of the polar bodies

A small number of previous publications were able to detect polar bodies (PBs) of stage 1 *P. tepidariorum* embryos at the periphery of the yolk, close to the vitellin membrane (Mittmann and Wolff 2012, Pechmann et al. 2017). To verify and repeat these observations, we fixed and marked the DNA (via SYTOXGreen staining) to visualize the polar bodies at stage 1 *P. tepidariorum* embryos. At this early stage of development, we could detect the polar bodies as a small peripheral cluster that was only present in one half of the embryos (see green arrowheads in Fig. 1 A and B).

**Fig. 1.**
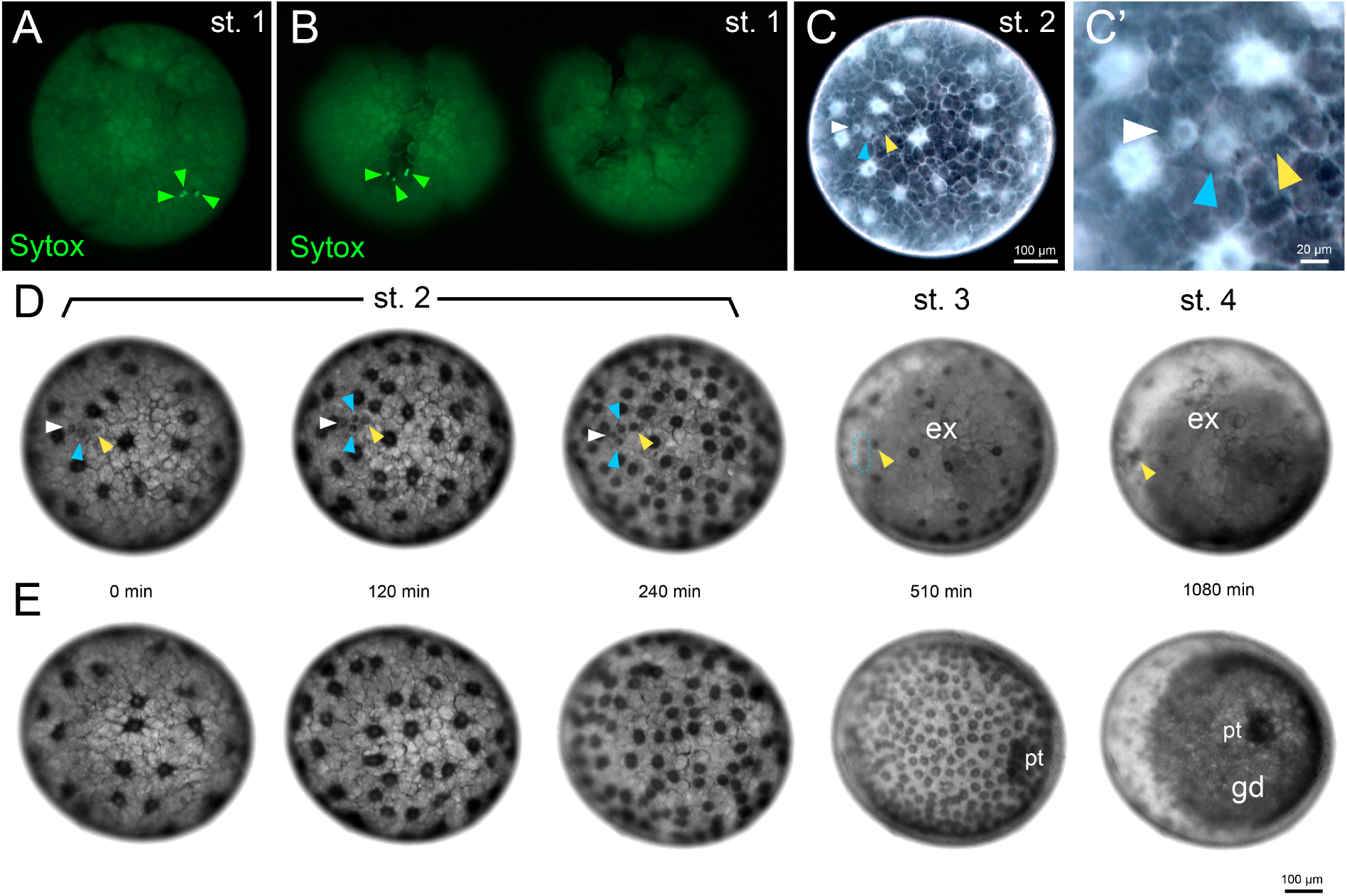
Polar body location during germ disc formation. (**A** and **B**) At early stage 1, the three polar bodies (marked by Sytox green and green arrow heads) are clustered at the periphery of *P. tepidariorum* eggs. (**B**) Two halves of a stage 1 *P. tepidariorum* egg. Only one half harbours polar bodies. A and B are formaldehyde fixed and Sytox green stained embryos. (**C-E**) Stills from Movie 1. Living *P. tepidariorum* embryos were imaged via transmitted light microscopy. Polar bodies are marked by white, blue and yellow arrow heads. A dividing polar body and resulting descendants are marked by the blue arrow heads (see D 0-240 min). (**D**) In this embryo, polar bodies became visible at stage 2 of embryonic development (see arrowheads at 0 min) and the germ disc formed opposite to the polar bodies (on the far side of the viewers perspective, compare to Movie 1). (**E**) No polar bodies were visible in this embryo and the germ disc did form on the near side of the viewers perspective (compare to Movie 1). An inverted image of the embryo shown in D (0 min) is shown in C. The region of the polar bodies is magnified in C’. To better visualize the polar bodies, the contrast was enhanced in C and C’. Abbreviations: pt – primary thickening; ex – extraembryonic; gd – germ disc.

In order to see whether the position of the polar bodies might correlate with the formation of the germ disc, we were searching for ways to visualize polar bodies in living embryos. By imaging and analysing the embryonic development of more than 400 embryos from various cocoons (see suppl. Fig. 1) we detected small cell-like structures that became visible at stage 2, when the cleaving energids did reach the cortex of the egg (see Fig. 1 C, C’ and D; e.g. Movie 1, 2, 7, 8). Like the cleaving energids, these small cell-like structures (see arrowheads in Fig. 1 C, C’ and D) possessed a small cytoplasmic island that surrounded a nucleus-like structure (see the white cytoplasm that surrounds the dark appearing nuclei in Fig. 1 C and C’). However, in comparison to the cleaving energids, these cell-like structures were smaller (around 20 μm vs. 40 μm, see scale bar in Fig. 1 C, C’) and did behave differently, in regard to their moving and splitting behaviour (see Movie 2). While the cleaving energids were dividing very synchronously during stage 2 (followed by a dynamic separation of the daughter energids), the small cell-like structures divided very irregularly and appeared more static in embryos that did not contract, yet. The number of these small cell-like structures and their clustered arrangement, (see Fig. 1 A and B), suggested that they correspond to polar bodies. As we were able to verify this assumption with further experiments (see below), we will name these small cell-like structures ‘polar bodies’ (PB), from now on.

Because of the very granular structure of the yolk, which led to a strong light refraction using transmitted light microscopy, the PBs were only hardly visible during stage 1 of embryogenesis (see Movie 2 and 7, suppl. Movie 1 and 6, Fig. 2 A).

**Fig. 2.**
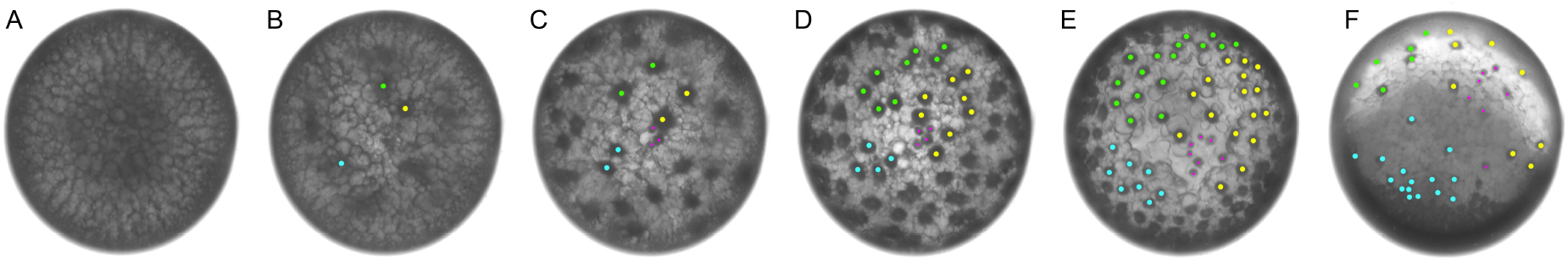
Polar bodies and polar body derived structures end up in the ab-embryonic hemisphere. **(A-F)** Stills from Movie 2. Extraembryonic energids (nuclei with surrounding cytoplasm) were tracked in cyan, green and yellow. Polar bodies (see **C**) and polar body derived structures (see **D-E**) are marked in magenta. (**F**) Polar body derived structures ended up at different positions within the extraembryonic hemisphere.

Now that we were in the position to see the PBs in living embryos we were able to analyse if there was a correlation between the position of the PBs and the forming germ disc. Indeed, by recording the formation of the germ disc of 412 *P. tepidariorum* embryos, we found that there was a clear tendency of the germ disc to form opposite to the PBs. As we were using transmitted light conditions and the yolk of spider embryos is very opaque, we failed to detect the PBs in most of the embryos. In a big portion of the analysed embryos (48.1%, see suppl. Fig. 1) the germ disc formed in a lateral position. As the transmitted light conditions in conjunction with the vitellin membrane and the chorion created a shadow at the “equator” of the embryo, we missed to detect the PBs in these embryos as the PBs were probably just at or a bit below the “equator” of the embryo. However, in more than one third of the embryos (37.3 %) we find a clear correlation between the position of the PBs and the formation of the germ disc. In 28.6 % of the analysed embryos (see suppl. Fig. 1), we detected the PBs in the embryonic hemispheres facing the camera. Interestingly, in these embryos the germ disc formed on the far side of the viewers perspective and the PBs ended up in the extraembryonic area (or contra-orbital region (Prpic and Pechmann 2022)) of the embryos. Additionally, in 8.7 % of the analysed embryos the germ disc did form towards the viewers perspective (see Fig. 1 E, suppl. Fig. 1; suppl. Movie 5). In these embryos, PBs were not detectable and were potentially on the side facing away from the camera.

6.3 % of the analysed embryos showed strong malformations and were not further taken into account (see suppl. Fig. 1 and suppl. Movie 8). In contrast, in 8.3 % of the recorded embryos we could detect some additional cell-like structures that appeared similar to the already described PBs but behaved differently. These structures divided rapidly and ended up in the germ disc of the embryos (see suppl. Fig. 1 and 2, Suppl. Movie 1, 2 and 7). As we could detect the PBs and additional cell-like structure in some of the analysed embryos (see suppl. Movie 1 and suppl. Fig. 2) at the same time, we assumed that these additional cell-like structures are neither PBs nor regular cleaving energids. At present, we must regard these additional cell-like structures as “unknown exceptions” that must be further analysed in future studies. Overall, this first analysis reveals a strong correlation between the localisation of the PBs and the germ disc forming opposite to the PBs.

### Dividing polar bodies

In the literature, polar bodies are often described as structures that degenerate over time (reviewed in e.g. Schmerler and Wessel 2011). Therefore, we were very surprised to detect PB divisions in our analysed spider embryos. Our tracking analysis clearly revealed that PBs divide (depicted in Figure 1 D and Figure 2, e.g. Movie 2) and up to seven PB derived structures could be detected at multiple regions within the extraembryonic region of stage 4 embryos (e.g. Fig. 2). At the beginning of stage 3, the contraction of the embryo is leading to an embryo that is freely floating within the perivitelline fluid. At the same time, germ disc condensation is leading to a redistribution of the yolk, which results in the extraembryonic region becoming heavier. This process led to an upward rotation of the germ disc making it impossible for us to track the PBs beyond stage 4 of embryonic development. In future studies, inverse microscopy could be used to track PB derived structures during later stages of development.

### Fluorescent live imaging of PB divisions and the process of germ disc formation

In order to better analyse the division pattern of the PBs and being able to detect PBs during stage 1 of embryogenesis we were looking for additional ways to detect PBs in living embryos. In 2022, Fujiwara and colleagues used several non-toxic fluorescent dyes to image and model germ band formation of *P. tepidariorum* embryos (Fujiwara et al. 2022). We used one of these dyes (SPY555-DNA) in our study and injected it to stage 1 *P. tepidariorum* embryos. This was in contrast to the study by Fujiwara and colleagues where the same dye was injected to stage 3 embryos, at the earliest. Although stage 1 injections did lead to a bright fluorescent spot at the injection site, SPY555-DNA was incorporated to the DNA of the polar bodies and we could detect three bright DNA spots of the PBs already 30 minutes after the injection of the dye (see arrowheads in Fig. 3 and the tracks in Fig. 4 A and A’; Movies 3 and 4). After injection of SPY555-DNA to *P. tepidariorum* stage 1 embryos we could detect the same behavior of the germ disc as in un-injected embryos, namely, the germ disc did form opposite to the polar bodies (Fig. 3 A; see the three embryos showing SPY555-DNA labelling of the PBs shown in Movie 3 B-D). In SPY555-DNA injected embryos that did not show any sign of PBs, the germ disc did form towards the viewers perspective, reflecting the results obtained for un-injected embryos (see Fig. 3 B and Movie 3 A; compare to Fig. 1 and Movie 1). Our experiments demonstrate that the formation of the germ disc is independent to the point of injection but is clearly related to the position of the PBs (compare the different positions of the PBs in the embryos shown in Movie 3 to the final position of the fully condensed germ disc).

**Fig. 3.**
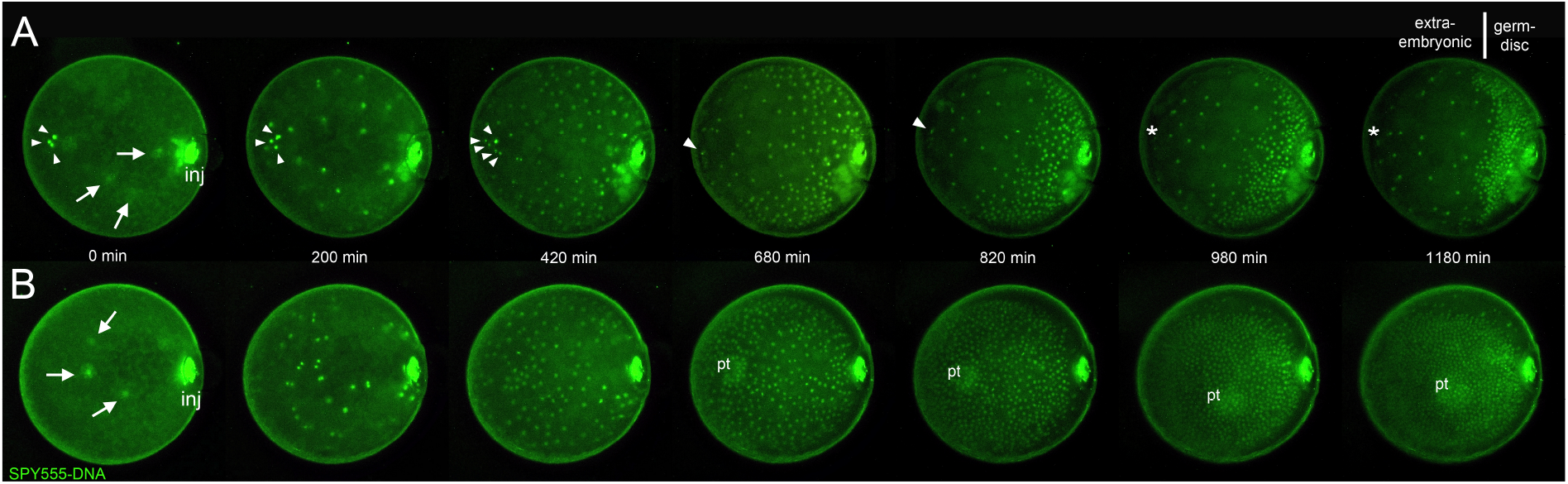
Fluorescent live imaging of polar bodies. (**A and B**) Stills of two of the four embryos shown in Movie 3. Fluorescent time-lapse recordings of living embryos injected with the live cell stain SPY555-DNA. Embryos were injected with SPY555-DNA at mid stage 1 of embryonic development. As excessive SPY555-DNA leaked from the injection site, the bright fluorescent injection site is always visible (see “inj” at 0 min in A and B). Time lapse recording was initiated around 1 hour after injection. (**A**) The DNA of the three polar bodies (arrowheads) as well as of the cleaving energids (see arrows in A and B at 0 min), showed a strongly fluorescent signal shortly after injection of SPY555-DNA. Dividing polar bodies are indicated by multiple arrow heads (see embryo in A at 420 min; compare to Fig. 4 A-C). As the contraction of the embryo (between 420 and 680 min, see Movie 3) is resulting in a slight rotation of the embryo, the DNA of only one polar body is still visible at later time points of development (see 680 and 820 min). As polar bodies disappeared completely from the field of view at 980 min, the former position of the polar bodies (opposite to the forming germ disc) is indicated by an asterisk. (**B**) No polar bodies were visible in the depicted embryo and the germ disc formed towards the viewers perspective. The forming primary thickening is indicated (see pt at 680-1180 min).

**Fig. 4.**
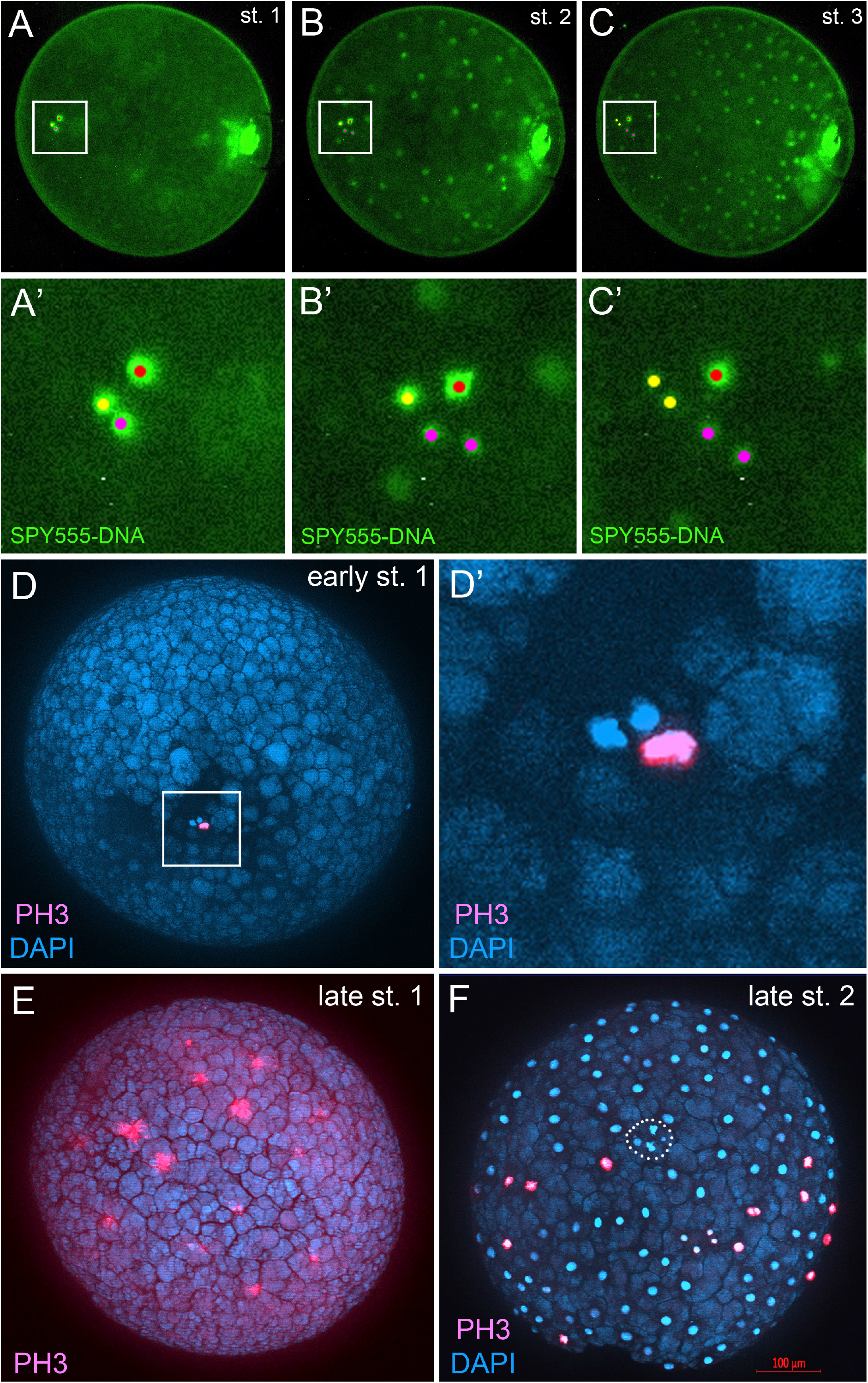
Dividing polar bodies. (**A-C**) Same embryo as shown in Fig. 3 A (see embryo in Movie 3 D). The DNA of the individual polar bodies was tracked until they disappeared from the field of view (see yellow, red and magenta dots, compare to arrowheads of Fig. 3A). (A’-C’) Magnified areas of the boxed regions in A-C. (**D-F)** Phospho-Histone H3 antibody (PH3) staining of stage 1 and 2 embryos. (**D and D’**) An early stage 1 embryo with a single polar body that is in the process of division. Cleaving energids are deep within the yolk and not yet visible in this embryo. (**E**) Synchronous energid cleavages of a late stage 1 embryo. Energids are about to reach the outer cortex region of the egg. (**F**) At late stage 2, cell divisions/energid cleavages are no longer synchronous. Potential polar bodies (see irregular and partially fragmented DNA) are encircled.

Interestingly, in the SPY555-DNA injected embryos, we find single embryos that show uncommon small, dividing cell-like structures that neither reflect PBs nor regular cleaving energids (see suppl. Movie 2). As we see similar structures in regular, un-injected wt embryos (see suppl. Movie 1), it is unlikely that these additional cell-like structures were induced by SPY555-DNA injections. As already mentioned above, the nature of these small cell-like structures is unclear and needs to be determined in future studies.

Injections of SPY555-DNA to stage 1 *P. tepidariorum* embryos followed by fluorescent time lapse recordings, clearly demonstrated that the nuclei of the polar bodies are dividing (see tracks in Fig. 4 A-C’ and Movie 3 and 4). In addition, the DNA (nuclei) of the polar bodies is (are) sometimes fragmenting/degenerating (see suppl. Fig. 3 and suppl. Movie 3). From our experiments it is unclear to what extend the DNA of the PBs is replicated and divided up between daughter PBs. An additional antibody staining against Phospho-Histone H3 (PH3) indicated single PBs that were entering division cycles at certain developmental time points (see Fig. 4 D and D’). This contrasts with the regular cleaving energids, that divide synchronous during stage 1 (see Fig. 4 E). At stage 4, when the germ disc was fully formed, the extraembryonic cells did no longer divide (see the missing PH3 signal in the extraembryonic region of the embryos shown in suppl. Fig. 4 B). However, at this stage we detected potential polar body derived DNA fragments in the extraembryonic area of the analyzed embryos (see arrowheads in suppl. Fig. 4 B).

Finally, we performed an *in-situ* hybridization with the gene *Delta*, which is strongly expressed in the cells of the extraembryonic region of the embryo (Oda et al. 2007). As the huge extraembryonic cells of *P. tepidariorum* embryos are rich in yolk granules (Pechmann 2016), transcripts are concentrated in the cytoplasmatic area that is close to the nuclei (see suppl. Fig. 4 A). Interestingly, we found some nuclei in the extraembryonic region that showed no *Delta* transcripts in their immediate vicinity (see arrowheads of suppl. Fig. 4 A). We assumed that these nuclei were derived from the polar bodies, which ended up in the extraembryonic area, but might not influence the patterning of the extraembryonic region.

### Conserved aspects of PB location and germ disc formation across spiders

*Parasteatoda tepidariorum* belongs to the cobweb spiders (Theridiidae), a species rich family (>2579 species in 129 genera) of the infraorder Araneomorphae, which make up the majority of the 52,338 spider species (World Spider Catalog 2024, see Wheeler et al. 2016 for spider phylogenies). While *P. tepidariorum* is a relatively distal branching cobweb spider, the genus *Steatoda* (closely related to black widows of the genus *Latrodectus*) branches basally within theridiid spiders (e.g. Liu et al. 2016, Wheeler et al. 2016). In Germany, *Steatoda grossa* is a frequently occurring species and a small culture of *S. grossa* was already established in our laboratory (see Wang et al. 2023). To confirm that the germ disc is not only forming opposite to the PBs in *P. tepidariorum* we analysed germ disc formation via time lapse recordings in *S. grossa* embryos. Indeed, also in *S. grossa* we could detect PBs that end up in the extraembryonic area after germ disc formation (Movie 5). This result shows the conserved aspect of germ disc formation opposite to the polar bodies within theridiid spiders.

The formation of a distinct germ disc was reported for many different spider species, including *Parasteatoda tepidariorum, Zygiella x-notata, Latrodectus mactans* and *Steatoda grossa* (e.g. Akiyama-Oda and Oda 2003, Chaw et al. 2007, Edgar et al. 2015, Wang et al. 2023). However, a distinct germ disc is not present in spider species like *Cupiennius salei* or *Cheiracanthium mildei* (Wolff and Hilbrant 2011, Edgar et al. 2015). Furthermore, as there seems to be no distinct germ disc in Mygalomorph spider species like *Ischnothele karschi*, and *Acanthoscurria geniculata* (reviewed in Edgar et al. 2015, Pechmann 2020) the formation of a distinct germ disc might be a derived feature. The difference between a distinct and non-distinct germ disc is due to the morphology and number of the extraembryonic cells. In spiders that show a distinct germ disc the number of extraembryonic cells is small and the cells have a squamous morphology. In contrast, the extraembryonic cells of spider embryos with a non-distinct germ disc are rich in number and the morphology of the extraembryonic cells is probably very similar to the cells that will give rise to the embryo. However, in spiders without a clear germ disc, a kind of boundary between extraembryonic and embryonic cells becomes eminent at a later stage of development (stage 5), when gastrulating cells spread only in the embryonic half of the egg (e.g. Pechmann 2020, reviewed in Prpic and Pechmann 2022). Regardless of the fact whether there is or is not a distinct germ disc present in different spider species, all spiders analysed so far, develop a centrally located primary thickening, during early (stage 3-4) spider development (e.g. Akiyama-Oda and Oda 2003, Wolff and Hilbrant 2011, Pechmann 2020, Holm 1954). The primary thickening is a source of gastrulating cells that will also lead to the formation of the migratory cells of the cumulus, which are important to pattern the DV body axis of the spider embryos (Pechmann 2020, Oda and Akiyama-Oda 2008, Oda et al. 2020). In spiders without a distinct germ disc, we wanted to see whether it would be possible to find a correlation between the polar bodies and the formation of the centre of the embryo, the primary thickening. In order of being able to determine conserved aspects we analysed the location of polar bodies and primary thickening formation in a basally branching spider, *Ischnothele caudata*, which belongs to the Mygalomorphae. In freshly laid *I. caudata* embryos, we could detect three SYTOX Green positive spots, which most likely represented the polar bodies (see Suppl. Fig. 5 A and B). Subsequently, we analysed living *I. caudata* embryos and performed time lapse recordings. In a number of *I. caudata* early blastoderm embryos, we could detect a small cluster of cells that most likely represented the polar bodies of this spider species (see Suppl. Fig. 5 C’ and Movie 6). As the extraembryonic cells divided quickly and massively reduced their cell surface, we could not track the polar bodies for a long time (see Movie 6). Nevertheless, our time lapse recordings demonstrate that in all *I. caudata* embryos where we could detect the potential PBs, the primary thickening formed on the far side of the viewers perspective.

In summary, our results indicate a conserved correlation between the position of spider polar bodies and the germ-disc/primary thickening forming on the opposite side to the PBs.

### Tracking and labelling of early cell clones across the spider embryo

The knowledge that the germ disc forms opposite to the polar bodies allowed us to track the fate of early cells of the blastoderm. Our tracking revealed that only the cells that are directly adjacent to the PBs ended up in the extraembryonic half of the embryo (see green cell markings at 32 and 64 nuclei stage in Fig. 5 A, Movie 7), while the next most distant cells already ended up at the rim after germ disc condensation (follow cell markings of the 128 nuclei stage up to stage 4 in Fig. 5 A, Movie 7) and will develop into anterior portions of the embryo (see McGregor et al. 2008 and Hemmi et al. 2018 for cell fates of the germ disc).

**Fig. 5.**
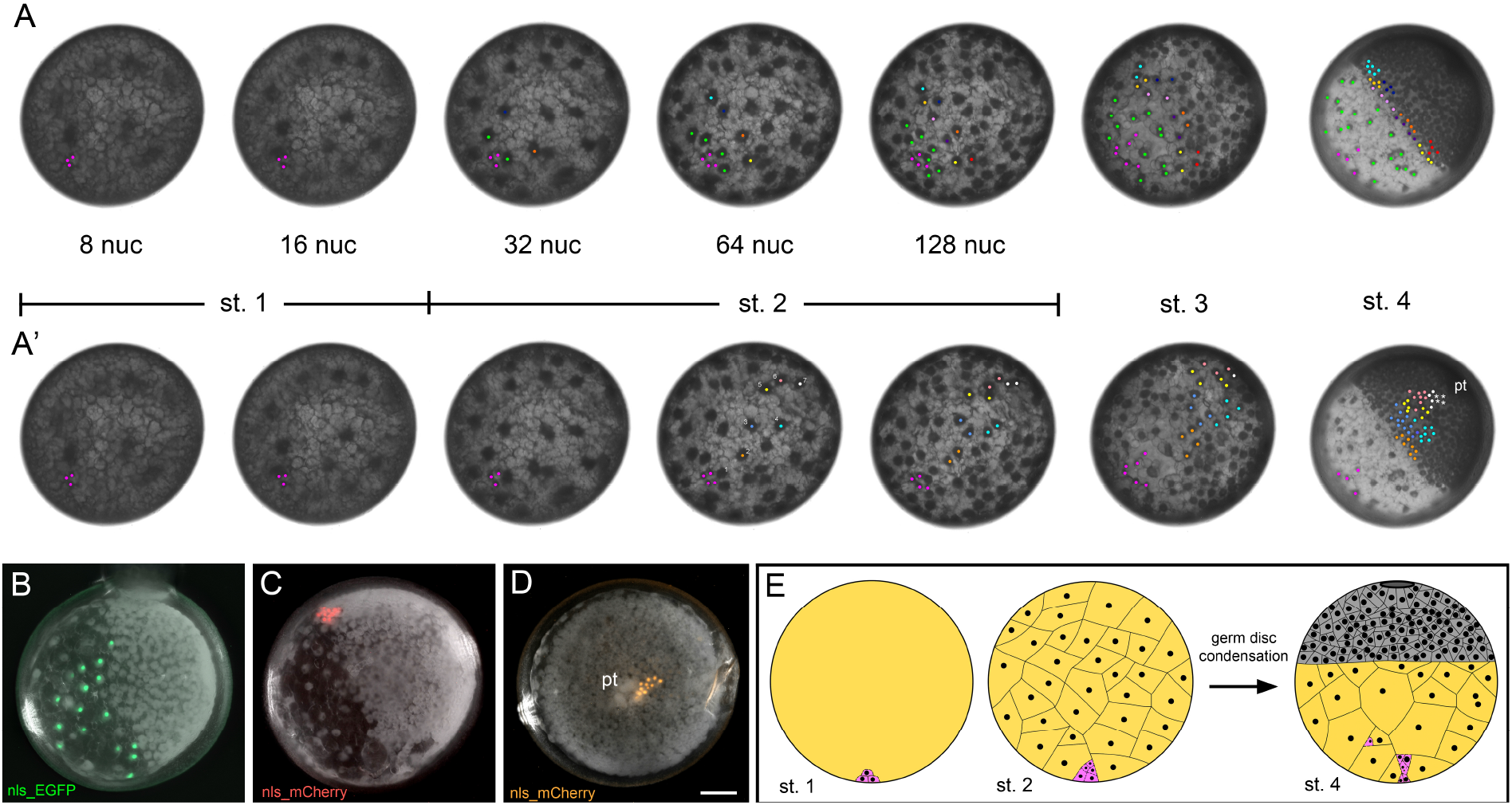
Polar bodies and applications. (**A and A’**) Stills from Movie 7 and 8. Polar bodies were tracked in magenta. (**A**) Extraembryonic cells were tracked in green. Additional tracking was carried out to track rim cells of the germ disc. (**A’**) Same embryo as shown in **A** but tracking a different subset of cells. Tracking starts at 64 nuclei stage to mark cell clones that end up at the periphery and more central regions of the germ disc at stage 4. Cells marked by the white asterisks indicate cells that most likely belong to the cell clone marked by the white dots. The numbering (1-7, see middle embryo in A’) indicates different distances of certain cells/energids with respect to the PBs (see text for more details). (**B**) At stage 2, capped mRNA coding for nuclear localized EGFP was injected to a cell that was directly next to the polar bodies. The resulting cell clone did spread over the extraembryonic area of the embryo. (**C**) Still from Movie 10 rotated by 180°. At the end of stage 2 (128 nuclei stage), capped mRNA coding for nuclear localized mCherry was injected 4 cells away from the polar bodies (see Movie 10). At stage 4, the resulting cell clone did end up at the rim of the germ-disc. (**D**) At stage 2, capped mRNA coding for nuclear localized mCherry was injected opposite to the polar bodies. The resulting cell clone ended up in close proximity to the primary thickening (pt). Scale bar is 100 μm. (**E**) Schematic summary of the findings presented in this study. Take home message: The spider germ disc forms opposite to the polar bodies. Polar bodies are marked in magenta, yolk is marked in yellow, yolk-free germ disc cells are indicated in grey, and nuclei are represented by black circles. The cytoplasmic island around the nuclei of the yolk rich blastoderm (at stage 2) and extraembryonic cells (at stage 4) has been omitted from the drawing. From stage 1 to stage 4, polar bodies often divide and sometimes disintegrate (compare to Movie 4, suppl. Movie 3 and suppl. Fig. 3; see tiny black dots at stage 4 within the centrally located polar bodies that indicate fragmented DNA). Finally (at stage 4), the polar bodies end up at different positions within the extraembryonic region of the embryo. The dashed lines between the polar bodies and the yolk rich extraembryonic cells (indicated in yellow) indicate a lack of knowledge of the exact morphology and integration of the polar bodies into the extraembryonic region.

Tracking different energids of a 64 nuclei stage embryo revealed that the energids/cells that were further away from the PBs could be found in more central locations of the disk, after the germ disc was fully formed (see Fig. 5 A’, Movie 8). With the polar bodies close to the “egg equator” but still in the field of view, we managed to track six energids/cells with different distances to the PBs (see the differently coloured energids at the 64 nuclei stage of Fig. 5 A’). During stage 2, the cleaving energids of *P. tepidariorum* embryos distribute in the form of strands (see Movie 7 and 8) and are never in a straight line. Therefore, we tracked the energids in a zigzag-like pattern (see orange to white coloured energids at 64 nuclei stage of Fig. 5 A’). Our tracking revealed that at 64 nuclei stage, an energid located seven positions away from the PBs (see zigzag numbering of the coloured energids in Fig. 5 A’) ended up next to the region of the primary thickening (see the energid/cell clone highlighted in white in Fig. 5 A’).

To verify our manual tracking results we injected capped mRNA, coding for nuclear localized fluorescent proteins, to single blastomeres of stage 2 embryos (Fig. 5 B-D, Movies 9 and 10). Injections directly next to the PBs resulted in extraembryonic cell clones, only (Fig. 5 B, Movie 9). In contrast, capped mRNA injections (coding for nuclear localized mCherry) at a 4 cell distance to the PBs (at 128 nuclei stage, see Movie 10) or directly opposite to the PBs resulted in mCherry positive cell clones that were either located at the rim of the germ disc (Movie 10, Fig. 5 C) or in central location of the germ disc close to the primary thickening of stage 4 embryos (Fig. 5 D). Overall, these result show once again that the germ disc/primary thickening forms opposite to the PBs and that only the energids close to the PBs end up in the extraembryonic area.

## Discussion and future directions

Our study identified the polar bodies as an early landmark for the AP body axis of spiders and demonstrates that the formation of the spider germ disc is not a stochastic but seems to be a predetermined process.

In most organisms PBs degenerate during the further maturation of the oocyte and in many animals the function of polar bodies is still poorly understood. However, in some animals, PBs seem to have a crucial function also during later developmental stages (reviewed in Schmerler and Wessel 2011). As examples, in scale insects and some parasitic wasps, PBs can either contribute to the bacteriome of the adult or they develop into extraembryonic membranes that surround the embryos (reviewed in Schmerler and Wessel 2011). Furthermore, in some parthenogenetic animals, the polar bodies fuse with the egg pronucleus to form a diploid cell that is than able to undergo embryogenesis (reviewed in Schmerler and Wessel 2011). Another interesting study on the fate of the polar bodies was identified by Sakai and colleagues using the moth *Bombyx mori* as a model system (Sakai et al. 2013). In *Bombyx mori*, the PBs seems to contribute the extraembryonic serosa (an important extraembryonic membrane that is required to protect the embryos against pathogens and desiccation (e.g. Panfilio 2008, Jacobs et al. 2013, Jacobs et al. 2014)), which develops opposite to the forming embryo. This situation would be very similar to the spider embryo, where the polar bodies also end up in the extraembryonic area (this study). It is unclear whether the polar bodies and polar body derived structures have any function during later stages of spider development and future studies are necessary to address this question.

PBs from as a by-product of female meiosis and in many animals the small PBs are “budding of” from the oocyte and are set aside. This is in contrast to arthropod embryos like *Drosophila melanogaster*, where the nuclei of the polar bodies stay at the periphery of the egg and are not extruded from the oocyte (e.g. Loppin et al. 2015). Also in spiders, the nuclei of the polar bodies seem to stay near the cortex of the oocyte (this study, Suzuki and Kondo 1994). Our DNA labelling, nuclei tracking and Phospho-Histine H3 antibody staining revealed that the polar bodies of the spider *P. tepidariorum* divided and sometimes degenerated during the early stages of embryonic development. However, the number of PB divisions seems to vary between embryos and it is unclear why and how PBs are able to divide. As polyspermy was suggested for the closely related spider *Achaearanea japonica* (Suzuki and Kondo 1994), dividing polar bodies might be the result of an additional fertilization process of single or multiple PBs when additional sperm enter the oocyte. This might also explain differences in the number of dividing PBs.

The observation that the germ disc forms opposite to the polar bodies will allow us to better investigate certain aspects of early spider development, in future studies. Instead of random blastomere injections, it will be possible to use the position of the polar bodies to precisely target different regions of the embryos. This will be especially important for the analysis of the extraembryonic area and of the cumulus, as these structures are formed by only a small number of cells/energids. The minority of energids/cells of a 32 or 64 nuclei stage embryo will contribute to the extraembryonic region of a stage 4 P. *tepidariorum* embryo (this study, Pechmann 2016). Depending on the distance from the polar bodies at 32, 64 and 128 nuclei stage it will also be possible to target different regions of the germ disc in a relatively precise manner (compare to Fig. 5 A’). In future studies, the location of the polar bodies will also be of great use to identify potential maternally localized factors that might be responsible for body axis patterning in spiders.

An important finding of our study is the detection of polar body-like structures in a basally branching mygalomorph spider. Our analysis revealed that also in embryos of *Ischnothele caudata* these polar body-like structures were opposite to the forming primary thickening. This observation reveals a potential evolutionary conservation of the process of germ disc/primary thickening formation opposite to the polar bodies. Future studies analysing the presence and the location of the polar bodies in different chelicerate species will show if axis formation processes across different chelicerate species are broadly conserved.

Overall, our analysis reveals a clear correlation between the position of the polar bodies and the formation of the germ disc/primary thickening in araneomorph as well as mygalomorph spider embryos. Future experiments are required to reveal whether the polar bodies are actively involved in the formation of the germ disc. As the placement of the germ disc/primary thickening is required to set the AP body axis in spiders, the polar bodies serve as a landmark for body axis formation in spiders.

## Acknowledgements

We would like to thank Siegfried Roth and Nikola-Michael Prpic-Schäper for commenting on the manuscript.

## Disclaimer

The funders had no role in study design, data collection and analysis, decision to publish or preparation of the manuscript.

## Authors’ contributions

M.P. designed the study, performed experiments and wrote the paper. R.W. performed experiments and commented on the manuscript. All authors gave final approval for publication.

