## Supplementary Information for "Polar bodies serve as a landmark for anteroposterior axis formation in spiders"

### Movie Legends

**Movie 1:** Time-lapse recording of two *Parasteatoda tepidariorum* embryos, starting at stage 2 of embryonic development. Polar bodies (marked in red) were only visible in the upper embryo. The complete process of germ disc formation is depicted. In the upper embryo the germ disc formed on the far side from the viewers perspective. As the yolk is semi-transparent the germ disc is visible as a shadow. Transmitted light conditions.

**Movie 2:** Time-lapse recording of a *P. tepidariorum* embryo, starting at stage 1 of embryonic development. Extraembryonic cells were tracked in cyan, yellow and green. In contrast to Figure 2, in which the polar bodies were tracked in magenta, polar bodies are not marked in this movie. This allows the clear identification of the polar body derived structures that spread within the extraembryonic area. Because of gravity, the embryo changed its orientation at the end of the movie. The movie stops at mid stage 5, when the cumulus has travelled half the distance to the rim of the germ disc. Transmitted light conditions.

**Movie 3:** Fluorescent time-lapse recordings of SPY555-DNA injected *P. tepidariorum* embryos. Embryos were injected with the live cell stain at late stage 1 of embryogenesis. Only the cleaving energids were visible in the embryo shown in A and the germ disc formed towards the viewers perspective. In the embryos shown in B-D three or four polar bodies/polar body derived structures were visible and the germ discs formed opposite to the polar bodies. After contraction of the embryos (around 800 min), polar bodies shifted their positions, as the embryos were freely floating within the perivitelline fluid (best visible in the embryo shown in C).

**Movie 4:** Same embryo as shown in Movie 3 D. Polar bodies and polar body derived structures were tracked in red, yellow and magenta. The division of two polar bodies was clearly detectable.

**Movie 5:** Time-lapse recording of *Steatoda grossa* embryos, starting at stage 1 of embryonic development. Polar bodies were tracked in magenta. In the five embryos in which polar bodies were visible, the germ disc formed opposite to the polar bodies. After contraction of the embryos (around 720 min), polar bodies shifted their position, as the embryos were freely floating within the perivitelline fluid. Transmitted light conditions.

**Movie 6:** Time-lapse recording of *Ischnothele caudata* (Mygalomorphae) embryos, starting at stage 2 of embryonic development. Polar bodies are boxed and marked in magenta. Polar body-like structures were visible in all four depicted embryos and the germ discs formed opposite to the polar bodies, on the far side from the viewers perspective. As the yolk is semi-transparent, the germ discs are visible as a shadow. Transmitted light conditions.

**Movie 7:** Tracking of polar bodies (magenta), extraembryonic cells/energids (green) and future rim cells of the germ disc (differently coloured) in an embryo of *P. tepidariorum*. Polar bodies became visible at mid stage 1, when the cleaving energids reached half the distance to the surface. The complete process of germ disc formation and the beginning of cumulus migration (early stage 5) is depicted. Because of gravity, the embryo changed its orientation at the end of the movie. Transmitted light conditions.

**Movie 8:** Same embryo as shown in Movie 7. Tracking of polar bodies (magenta) and future germ disc cells (differently coloured) in an embryo of *P. tepidariorum*. Depending on the distance to the polar bodies, tracked cells ended up in ever more central areas of the germ disc.

**Movie 9:** An embryo of *P. tepidariorum* in which a single cell (at 128 nuclei stage) was injected with capped mRNA coding for nuclear localized EGFP. Injection was close to the polar bodies (marked in magenta; PBs). The EGFP labelled cell clone divided, and daughter cells ended up in the extraembryonic area. The germ disc formed on the far side of the viewers perspective. Brightfield condition with fluorescent overlay.

**Movie 10:** An embryo of *P. tepidariorum* in which cells (at 128 nuclei stage) were injected with capped mRNA coding for nuclear localized mCherry. At least two adjacent cells were labelled at the same time. Polar body derived structures are marked in magenta. From the position of the injected cells towards the polar body derived structures, a gap of at least 3 unlabelled cells was visible. The mCherry labelled cell clones further divided and daughter cells ended up at the rim of the germ disc. Brightfield condition with fluorescent overlay.

**Supplementary Movie 1:** Time-lapse recording of an embryo of *P. tepidariorum* that showed a cluster of undefined cell-like structures (marked in red) at mid stage 1 of development. Shortly afterwards, the polar bodies became visible (marked in magenta) at a different position within the early stage 2 embryo. The germ disc formed opposite to the polar bodies and the cell-like cluster likely ended up within the germ disc (see dark patch at 12 o'clock location within the germ disc). Also compare to a similar cluster of cell-like structures that ended up in the germ disc of the embryo shown in Supplementary Movie 7. The nature of these additional cell-like structures is unknow, so far. Transmitted light conditions.

**Supplementary Movie 2:** Time-lapse recording of an embryo of *P. tepidariorum* that showed a cluster of undefined cell-like structures that rapidly divided and ended up in the germ disc. The embryo was injected with the live cell stain SPY555-DNA. The movie shows the same embryo in the fluorescent (left) as well as in the bright field (right) channel. As the germ disc forms towards the viewers perspective the polar bodies were potentially located on the far side of the viewers perspective and could not be detected via this imaging approach. The nature of these additional cell-like structures is unknow, so far.

**Supplementary Movie 3:** Fluorescent time-lapse recordings of a SPY555-DNA injected *P. tepidariorum* embryo. One of the polar bodies already divided (four polar bodies are present at the beginning of the movie) and the DNA of an additional polar body started to fragment at minute 520. A closeup of this fragmentation process is depicted.

**Supplementary Movies 4-8:** Supplementary Movies 4-8 represent the time lapse recordings of the embryos shown in Supplementary Figure 1. These embryos were chosen as a representative for each of the depicted categories (I-V) that were included into the statistical analysis.

### Supplementary Figures

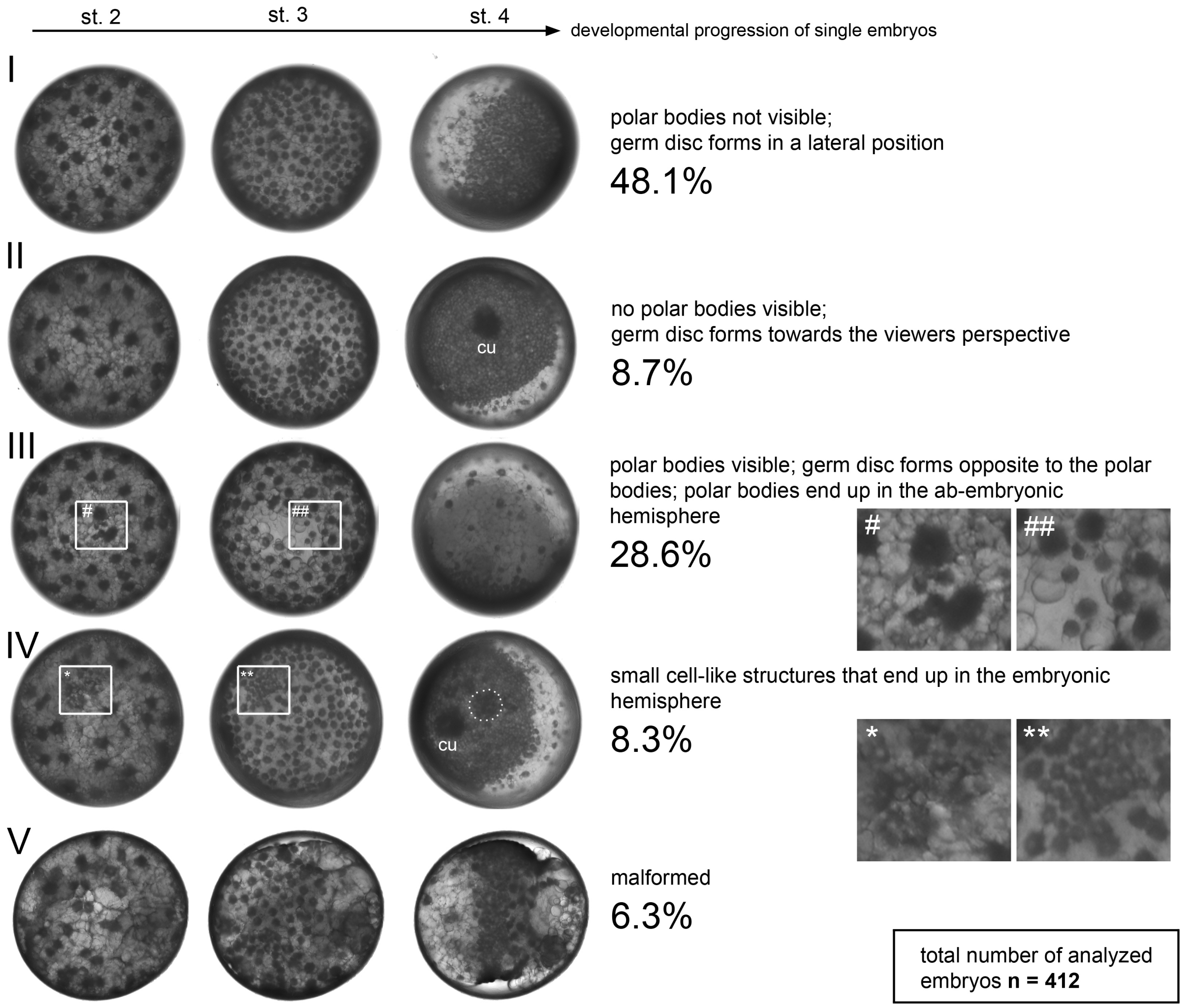


**Supplementary Figure 1.** Statistical analysis. Correlation between the location of the polar bodies (PBs) and the final location of the germ disc. The development of 412 embryos of two different cocoons from two *P. tepidariorum* females, were recorded via time-lapse imaging. Embryos were manually analysed and sorted into five (I-V) different categories. Each row shows one representative embryo of each category at three different developmental stages (stages 2-4, see Supplementary Movies 4-8). Insets marked by #, ##, * and ** are shown in a magnified view. The cumulus (cu) is marked in some embryos. The dashed line (see IV st. 4) indicates the region in which the cluster of the small cell-like structures did end up after germ disc condensation (compare to Suppl. Movie 7).

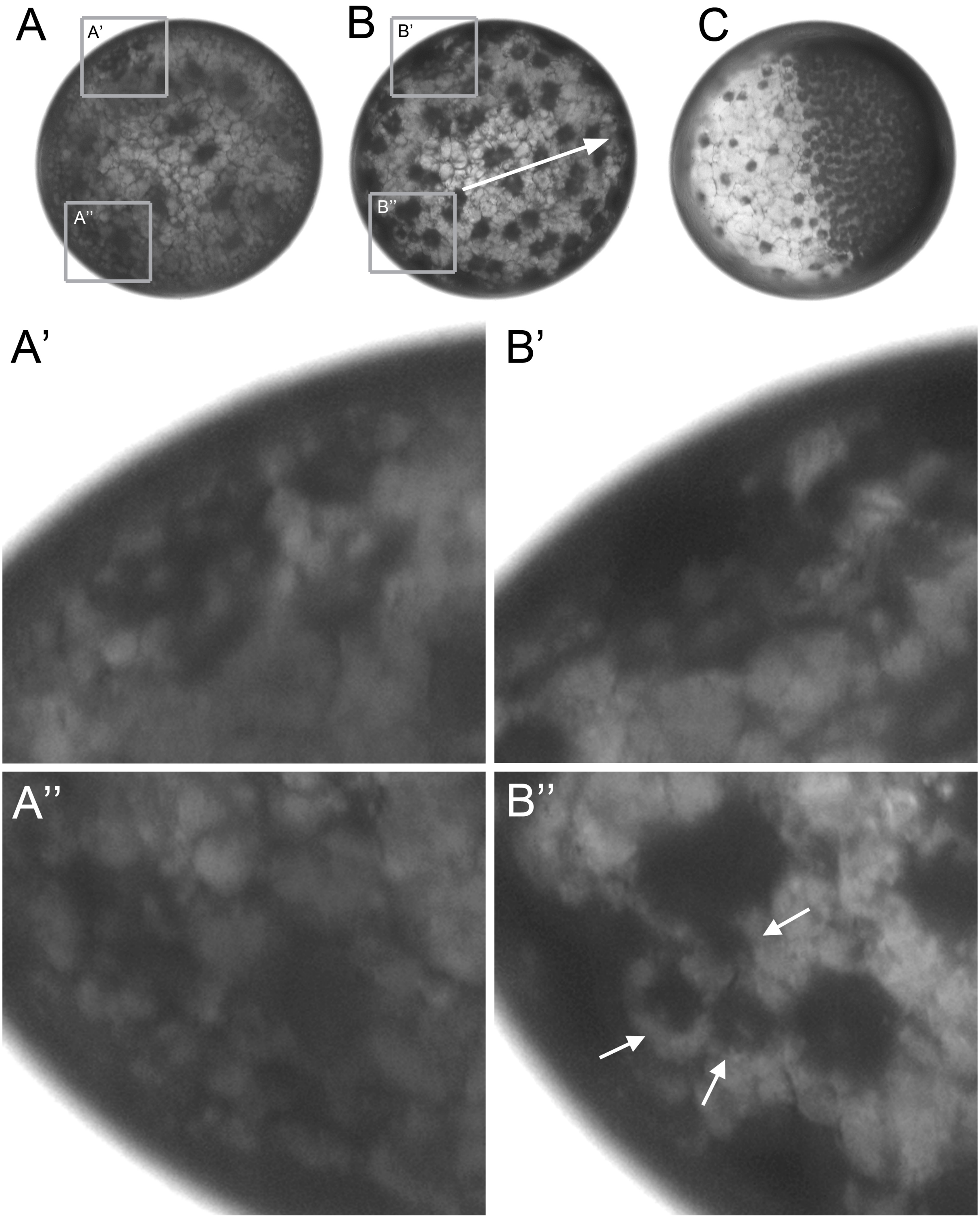


**Supplementary Figure 2.** (**A-C**) Stills from Supplementary Movie 1. Same embryo at late stage 1 (A), stage 2 (B) and stage 4 (C). Polar bodies are hardly visible at late stage 1 (see A’’). The polar bodies (see three white arrows in B’’) as well as some undefined cell-like structures (magnified in A’ and B’; see category IV in Supplementary Figure 1) were visible in the depicted embryo. The germ disc formed opposite to the polar bodies. The white arrow in B indicates the direction of the germ disc condensation process in respect to the position of the PBs. The undefined cell-like structures show no correlation with regard to the final position of the germ disc and likely end up in the germ disc (see figure legend of Supplementary Movie 1).

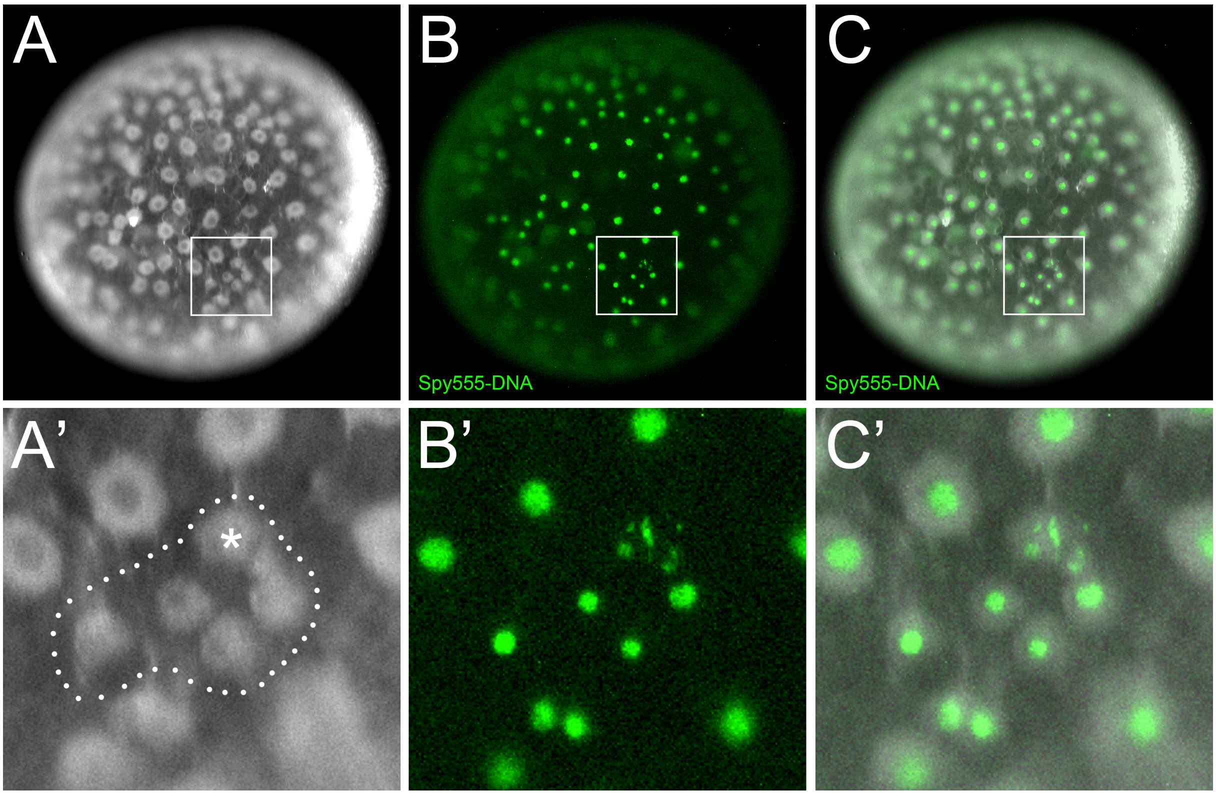


**Supplementary Figure 3.** Degenerating polar bodies. (**A-C**) An early stage 3 embryo that was injected with the live cell stain Spy555-DNA. Potential polar bodies are encircled (see dots in A'). The polar body that showed fragmented DNA (see B' and C') is indicated by the white asterisk in A'.

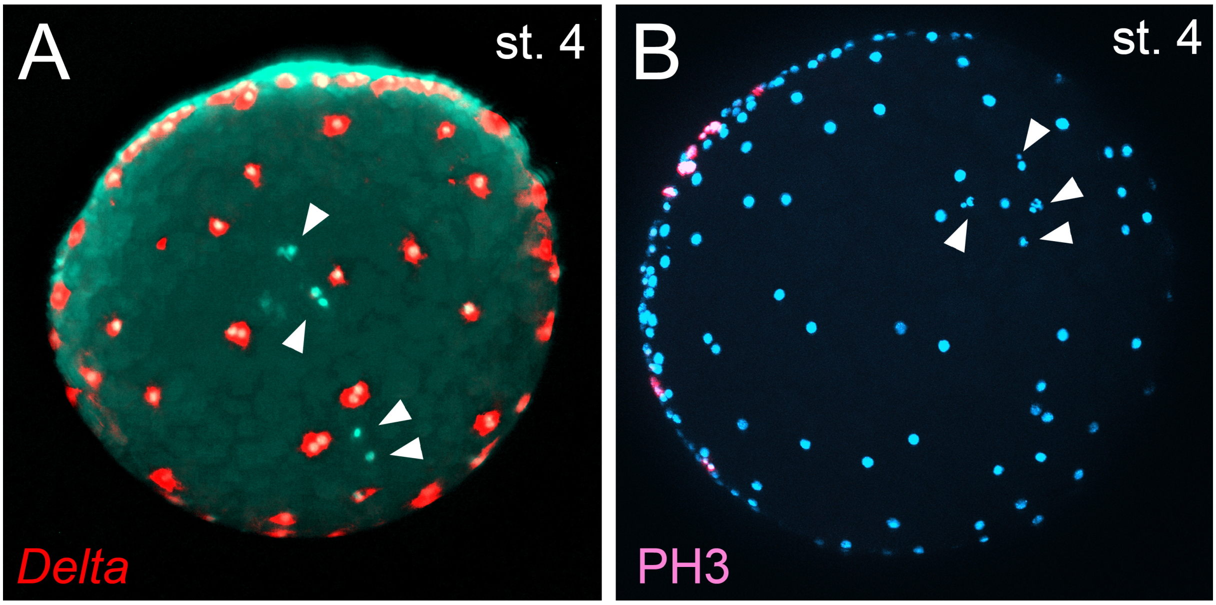


**Supplementary Figure 4.** (**A**) *In situ* hybridisation (false-colour overlay) detecting *Delta* transcripts. Depicted is only the extraembryonic area of a stage 4 *P. tepidariorum* embryo. *Delta* is strongly expressed in extraembryonic cells. Potential polar body derived structures (see arrowheads) were negative for *Delta* transcripts. (**B**) Phospho-Histone 3 antibody staining of a stage 4 embryo. Depicted is only the extraembryonic area of a stage 4 *P. tepidariorum* embryo. Potential polar body derived DNA fragments are indicated by the arrowheads. At stage 4, cell divisions were restricted to the germ disc cells.

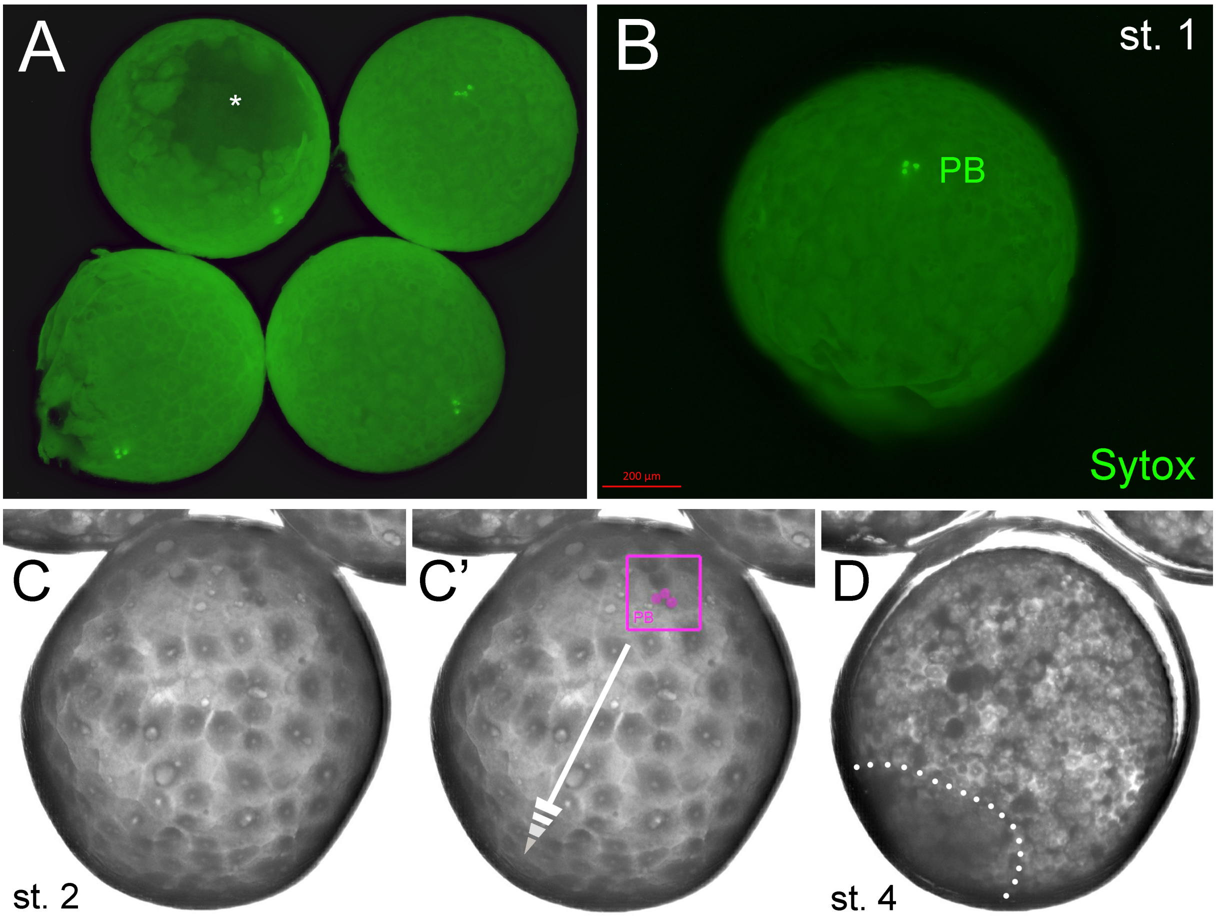


**Supplementary Figure 5.** Polar bodies in embryos of the mygalomorph spider *Ischnothele caudata*. (**A**) Four stage 1 embryos of *I. caudata* stained with SytoxGreen. Polar bodies are clearly visible. Stage 1 embryos are very fragile and some embryos were partially damaged during the devitellinization process (see asterisk) (**B**) A single stage 1 *I. caudata* embryo (stained with SytoxGreen) in a magnified view. The DNA of three polar bodies is clearly visible. (**C-D**) Stills from Movie 6. (**C**) Early blastoderm (st. 2) embryo of *I. caudata.* Potential polar bodies (PB) are marked in magenta in C'. The primary thickening formed opposite to the polar bodies and was visible as a shadow (indicated by white dots) through the yolk. In C', the direction of primary thickening formation (opposite to the PBs) is indicated by the arrow. The dashed arrow indicates that the primary thickening formed on the far side of the viewers perspective.
